# The impact of collective load transport on the individual walking

**DOI:** 10.1101/2023.11.17.567612

**Authors:** N. Sghaier, C. Pasquaretta, N A. Turpin, P. Moretto

## Abstract

Collective handling is a collaborative strategy that involves two or more people in carrying out load transport. Different positions can be adopted, depending on the handle locations of the transported load, external constraints and the capacities of the carriers. The most adopted collective transport in our daily life is stretcher type transport. However, very little research has focused on the kinematic modifications caused by this type of transport. This research aims to evaluate and quantify the modifications of the locomotor pattern of stretcher transport. Our results highlighted a modification of walking parameters (step length, duration of the walking cycle, speed of execution, etc.), an increase in energy cost but above all a modification of the walking pattern with a reduction in joint range of motion. These results could be used to establish new recommendations for musculoskeletal disorders.

## 1. Introduction

Team lifting is a collaborative strategy that involves two persons or more in the transport of a load. Various configuration can be found in real life situations, with various placements of the hand on the object, various external constraints (e.g., weight of the object, corridors), and various capacities and preferences of the carriers (Barrett & Dennis, 2005). These parameters were studied to some extent in the scientific literature, although mainly for singles carriers. For example, researchers studied the optimal boxes handle positions during individual manual handling that could increase user satisfaction (Jung & Jung, 2010). Load characteristics such as shape and dimension were shown to play an important role in the ability to comfortably carrying a weight (Garg & Saxena, 1980). In 1981, The National Institute for Occupational Safety and Health (NIOSH) established an ergonomic equation evaluating weight limits for lifting tasks for single individuals which take into consideration the vertical location of the load, the distance the load is lifted, the frequency of lifting and, the symmetric or non-symmetric aspect of the lifting. However, there is a lack of information for providing clear instructions about how to lift a load in a collective configuration.

So far, mainly individual liftings have been studied (Datta & Ramanathan, 1971; Heglund et al., 1995). According to these researches, the best way to carry a load individually in terms of ergonomy is to use the double pack mode and the head mode. In contrast, carrying the load by hand was the worst method to transport a load. These data mainly suggested that the load should be as aligned as possible with the center of mass of the subjects to avoid creating a destabilizing torque. However, the ergonomic evaluations were only based on physiological parameters such as oxygen consumption and heart rate.

The collective aspect of lifting has received much less attention and is more often studied for practical/applied cases such as observed in health care facilities (Haiduven, 2003; Barrett & Dennis, 2005), construction workers (Faber et al., 2012; van der Molen et al., 2012; Anton, Mizner & Hess, 2013), rescue activities (Gamble et al., 1991) or military (Sharp et al., 1997; Knapik, Reynolds & Harman, 2004). Transporting a casualty on a stretcher is commonly studied by research in order to evaluate individual performance and hand grip strength recovery (Knapik, Harper & Crowell, 1999; Leyk et al., 2006, 2007). Armstrong et al. (2020), ranked paramedic lifting task using a measure of biomechanical exposure and showed that the worst activity while working was lifting a scoop board from the ground to the waist. Moreover, they showed that the position of the carrier, head or foot end of the equipment, didn’t modify the biomechanical exposure.

Few biomechanically based investigations of collective stretcher transport have been conducted. Lanini et al. (2017) studied human walking gait kinematics adaptations during load transport and considered it as a quadrupedal gait. The results showed an overall modification in gait parameters such as step length, gait cycle time and, Center of Mass vertical displacement associated to a walking gait synchronization of the participant. Sensory and tactile feedback in human walking helps dyads to synchronize their movements during side-by-side walking (Nessler & Gilliland, 2009; Zivotofsky, Gruendlinger & Hausdorff, 2012; Sylos-Labini et al., 2018; Felsberg & Rhea, 2021). Fumery et al. (2018), studied the walking efficiency during a side-by-side collective load transport and, showed the ability of humans to collaborate efficiently during load carriage. However, all of these researches focused more on the collective modifications during a load transport and not on how the collective tasks impacted the individual performance. This collective movement could be mainly performed in two ways: (1) the team members look at each other and (2) they look at the same direction to transport a load. The second technique implies a forward and a backward walking of one team member, which could play a major role in the individual and collective performance of the load transport. For now, most of the researches described the backward walking pattern as a simple kinematic time reversal of the forward walking pattern and reported only few differences (Thorstensson, 1986; Winter, Pluck & Yang, 1989; Lee et al., 2013). However, backward walking in a collective transport task has not been investigated.

In this research we looked forward to explore different aspects of a collective stretcher, and mainly how does this kind of transport impact the biomechanical individual behavior. The objective of this research is to address three scientific questions: (1) What are the differences across forward and backward walking? (2) How a collective load transport impacts the biomechanics of an individual forward walking? and, (3) Does the backward walking play a major role in the biomechanical modification during collective load transport?

## 2. Materiels and methods

### 1.1. Population

Twelve women and eight men were recruited from September 01, 2019, to July 01, 2022. All participants were free of musculoskeletal or neurological disorder that might have affected the carriage. Mean (±s.d.) age, height and body weight were 24 ± 2,6 years, 1,71 ± 0,07 cm and 64,65 ± 8 kg, respectively. Each duo was matched by gender, height and weight. The project was approved by the University Review Board and all participants gave their written and, oral consent in accordance with the Helsinki convention.

### 1.2. Experimental

In total, 10 dyads performed four different lifting conditions. For the two first conditions, we instructed the participants to walk individually at their own pace in a Forward direction “C1” and, in a Backward direction “C2”. Then two collective conditions consisted in transporting collectively a stretcher shaped load of 1.2 kg with the participant 1 at the front and the participant 2 at the back (Figure 1.c). We instructed the participants to transport the stretcher shaped load while looking at the same direction (Figure 1.a); meaning that both participants were performing a forward walking “C3”. And we instructed them to perform the same condition while looking at each other (Figure 1.b); meaning that the participant 1 was performing a backward walking and, the participant 2 was performing a forward walking “C4”. All conditions were randomly presented. Three trials were recorded in each condition.

**Figure 1:**
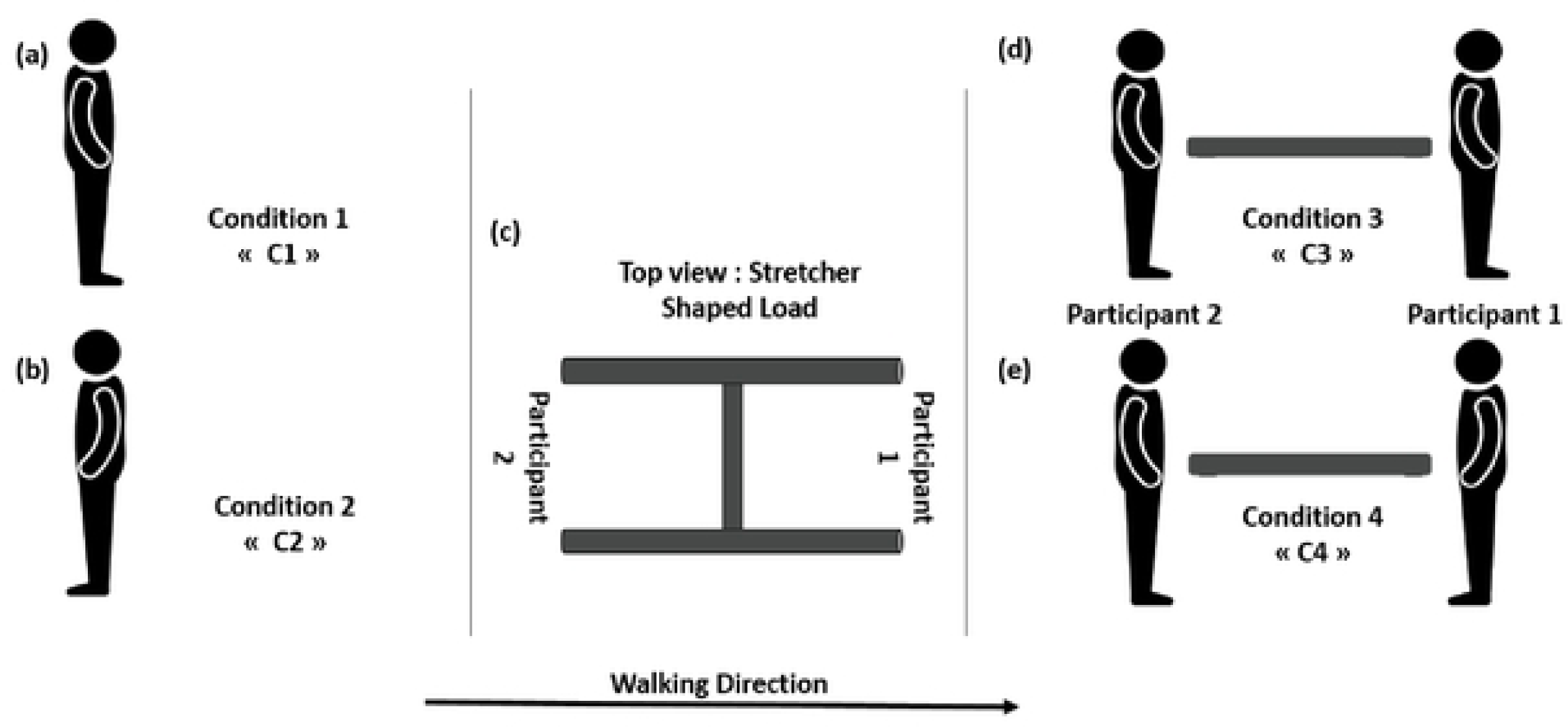
Experimental protocol with the placement of the participants for the condition 1 (a), condition 2 (b), condition 3 (d) and, condition 4 (e). With (c) representing the top view of the stretcher shaped load.

### 1.3. Kinematic data acquisition

Motion capture data were collected using 20 infrareds (11 Vero V2.2, 7 MX3 and, 2 MX TS40) transmitter-receiver video cameras (Vicon, Oxford metric’s, Oxford, United Kingdom) sampled at 200 Hz. Forty-two retro-reflective markers were placed on bony landmarks and on the navel of each participant (Wu *et al*., 2002, 2005), and 13 on the stretcher shaped load. One gait cycle per trial and participant has been considered for the analysis. These cycles were selected close to the middle of the travel path to avoid the acceleration and deceleration phases. The gait cycle of walking is defined by two successive foot strikes of the same foot. Regarding the collective transport C3 and C4, we investigated the Poly-Articulated Collective System (PACS) gait cycle, formed by the two participants and the load they carry, and defined by the first heel strike of the participant 2 (at the back) and the second foot strike of the participant 1 (at the front). The three-dimensional reconstruction of the markers position was performed using the Vicon Nexus 2.11.0 software.

### 1.4. Computed parameters

#### 1.4.1. Gait parameters

**Step length** was computed as the distance travelled between the foot strike of one foot and the strike of the contralateral foot.

**Gait Cycle Time** was computed as the time between two consecutive foot strikes of the same foot.

#### 1.4.2. CoM trajectory

De Leva anthropometric table (de Leva., 1996) was used to estimate the masses of each segment (*m_i_),* the position of the center of mass of each segment *i* (*CoM_i_*) of the participants and to determine the global position of the CoM of each carriers (i.e. CoM_Participant1_, CoM_Participant2_) and of the whole Poly-Articulated Collective System (PACS) which included the two subjects and the carried load (CoM_PACS_). The participants and PACS CoM location were all computed in the global frame of reference R(0, x, y, z) as follows (*m*_Participant1/2_, Participant 1 or 2 mass; *m*_PACS_, PACS mass):

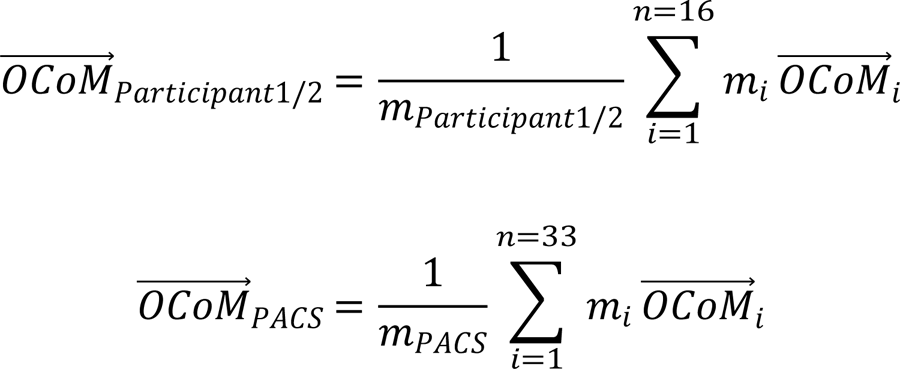

We then assessed the vertical amplitude (A = Zmax – Zmin, with Z the CoM height), the CoM period (P, which refers to the time between two peaks of the z-position curve) and, the CoM Velocity (Winter (1995).

#### 1.4.3. Recovery rate

The Recovery Rate (RR) assesses the amount of energy transferred between the potential and the kinetic energy of the center of mass (Bastien *et al*., 2016; Fumery *et al*., 2018a; Sghaier *et al*., 2022). RR is related to the consistency of the locomotor pattern and is based on the analysis of an inverted pendulum system (IPS) (Cavagna, Saibene & Margaria, 1963; Willems, Cavagna & Heglund, 1995; Gomeñuka *et al*., 2014). RRà was computed for each participant and the PACS as follow:

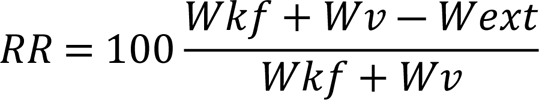

With Wkf the forward kinetic, Wv the vertical work and Wext the external work. These parameters were computed according to the method of Bastien *et al*. (2016).

#### 1.4.4. Joint angles

In order to study the kinematic modifications during a walking and a transporting task, we computed the joint angles at the hip, knee and ankle joints during a complete gait cycle. These joint kinematic were calculated according to the method proposed by the “Standardization and Terminology Committee” of the ISB, which defined a set of coordinate systems for various joints of the lower body based on the joint coordinate system and the XYZ sequence (Wu & Cavanagh, 1995; Wu *et al*., 2002).

The flexion/extension joint angles of the hip, knee and ankle were computed for each participant and condition, in order to highlight gait modification. According to Winter *et al*., (1989), the angles to time curve has been reversed from 100% to 0% of the gait cycle to compare backward to forward walking.

### 1.5. Statistical Analysis

Statistical analysis was performed with the software R v4.0.2 (http://www.r-project.org/). We compared the CoM velocity, step length, GCT, RR, CoM period and CoM amplitude across the four experimental conditions (C1, C2, C3 and, C4) using linear mixed models approach. We then estimated marginal means for each condition using the package *emmeans* in R (Searle, Speed & Milliken, 1980; Lenth *et al*., 2023; https://CRAN.R-project.org/package=emmeans). In order to analyze joint angle evolution, we used the Laassel *et al*. (1992) method. The gait cycle was divided into 25 windows, each of 4% of the gait time duration. To evaluate the effect of backward walking on the individual and collective walking, we ran 6 different linear models for each participant (P1 and P2) and each joints (hip, knee and ankle). Each model was built using conditions, windows and their interaction as predictors and angles as response variable. We then estimated the angle marginal means and their confidence interval for each condition in each window using the package *emmeans* in R.

## 3. Results

### 1.6. Gait parameters

The step length of participant 1 and 2 where significantly longer for C1 compared to C2 (respectively by +20.5% and +26.1%), C3 (respectively by 7.8% and 9.1%) and C4 (respectively by 19.6% and 14.9%). These differences were of +16% between C2 and C3 (C2>C3) and +12.8% between C3 and C4 (Figure 2.a). As for the second participant, its step lengths during C2 were significantly shorter than those performed during C1 (−35.3%), C3 (−22.9%) and C4 (−15.1%).

**Figure 2:**
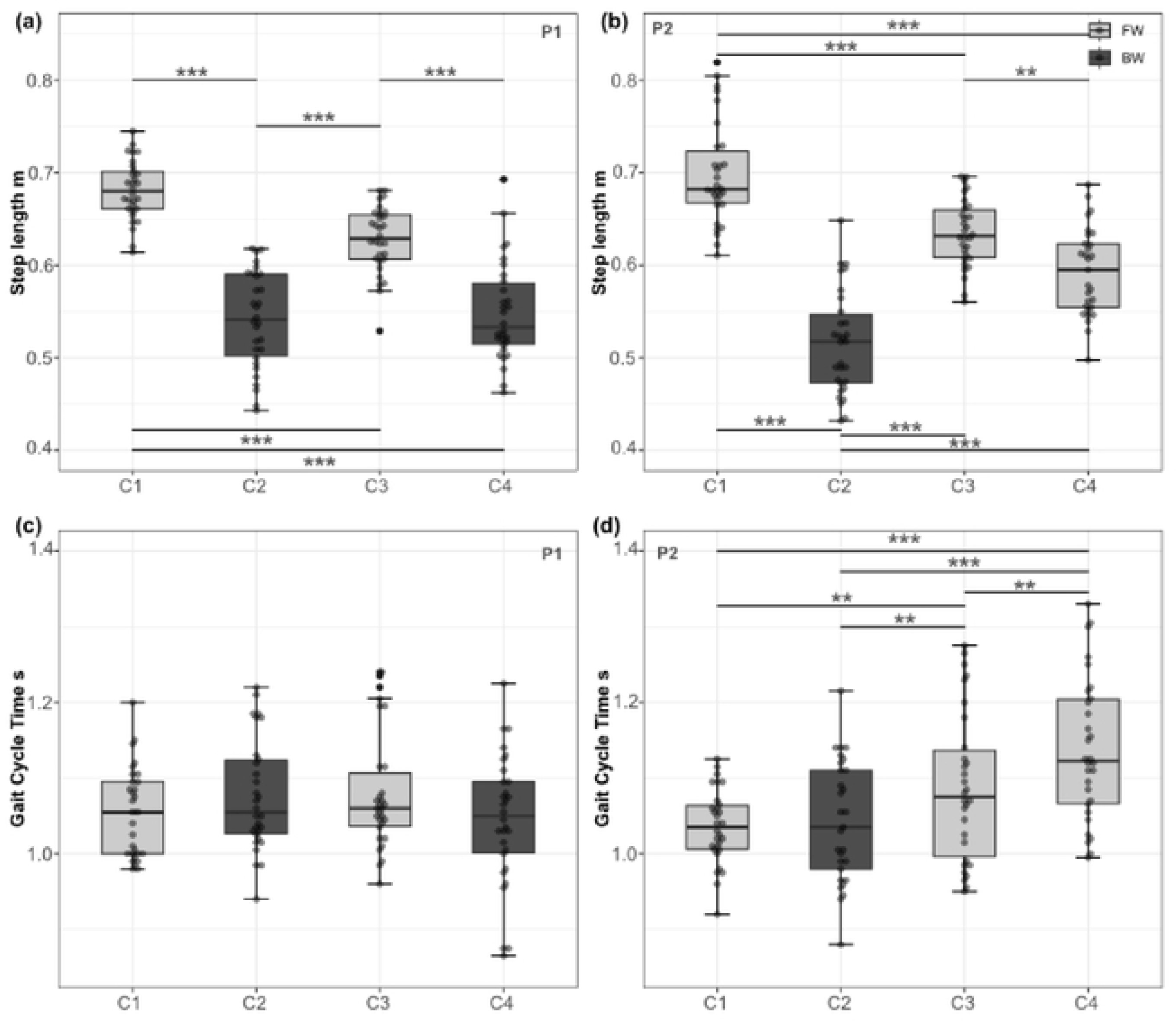
Step length and Gait Cycle time boxplot for participant 1 respectively (a), (c) and participant 2 (b) and (d) for condition 1,2, 3 and, 4. Light grey boxplots represent the forward walking performance and, dark grey boxplots represent the backward walking performance(* p< 0,05; ** p<0,01 and; *** p<0,001).

Globally for the GCT, there were no significant differences for the participant 1 between the different conditions. However, Participant 2 GCT was 7% greater in the collective conditions (C3, C4) than in the individual conditions (C1, C2). In addition, the GCT of C4 was 4.3% longer than the one of C3 (Figure 2.d).

### 1.7. Center of Mass (CoM) excursion

In order to compare the evolution of the CoM vertical excursion during a gait cycle (Figure 3), we computed the mean CoM amplitude and the CoM period for each trial (Figure 3). Globally, no differences were found for the individual CoM vertical excursion of P1 or P2 between the different conditions. However, the CoM amplitude was 17.2% higher during forward walking C1 than during backward walking C2. In addition, PACS’s CoM excursion was also 45.2% significantly higher in C3 than in C4. Concerning the CoM period, all conditions were the same whether they were performed individually or collectively.

**Figure 3:**
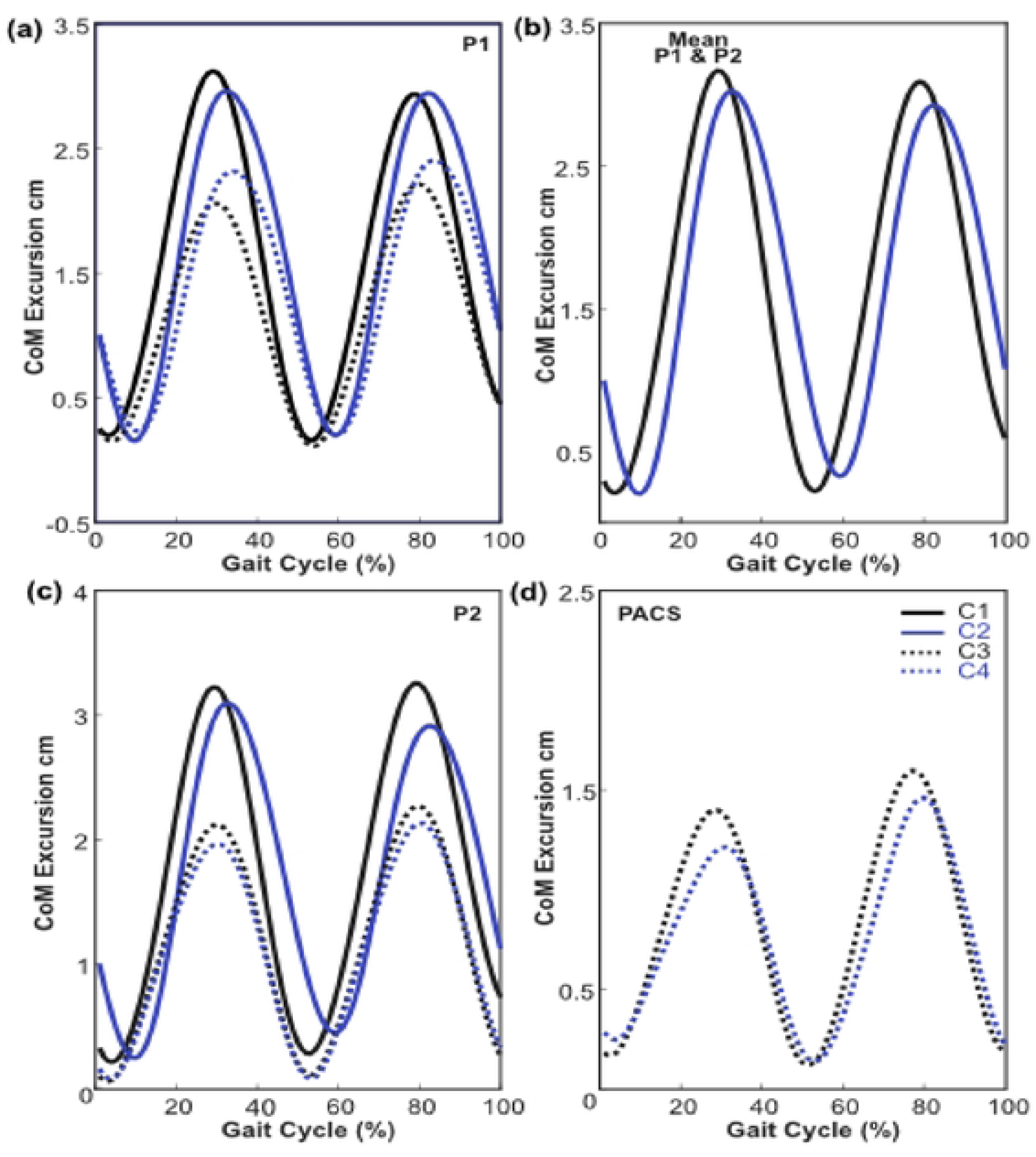
Center of Mass mean vertical excursion (10^-2^) for participant 1 (a) and (b), participant 2 (b) and (c) and, PACS (d). Continous lines represent the individual conditions C1 and C2 and, the dashed lines represent the collective conditions C3 and C4. The black line correpond to the forward walking performance and, the blue ones to the backward walking performance.

### 1.8. CoM velocity

For P1 the velocity performed during the first condition of individual walking decreased significantly for C2, C3 and, C4 (respectively by 21.6%, 10.9% and, 19.3%), which is also the case for P2 (respectively by 24.6%, 11.9% and, 21%) (Figure 4.a,b). Forward walking appears always faster than backward walking whether the participants are linked or not. Indeed, P1 and P2 increased their walking speed by 13.6% and 16.7% from C2 to C3. This is confirmed from C3 to C4 with a mean decrease of 10% for P1 and P2. Individually, backward walking induces a 23% decrease of the velocity (Figure 4.c). It is confirmed when the participants carry the load. The velocity of the PACS during C3 decreased by 9.9% compared to the one computed for C4 (Figure 4.d).

**Figure 4:**
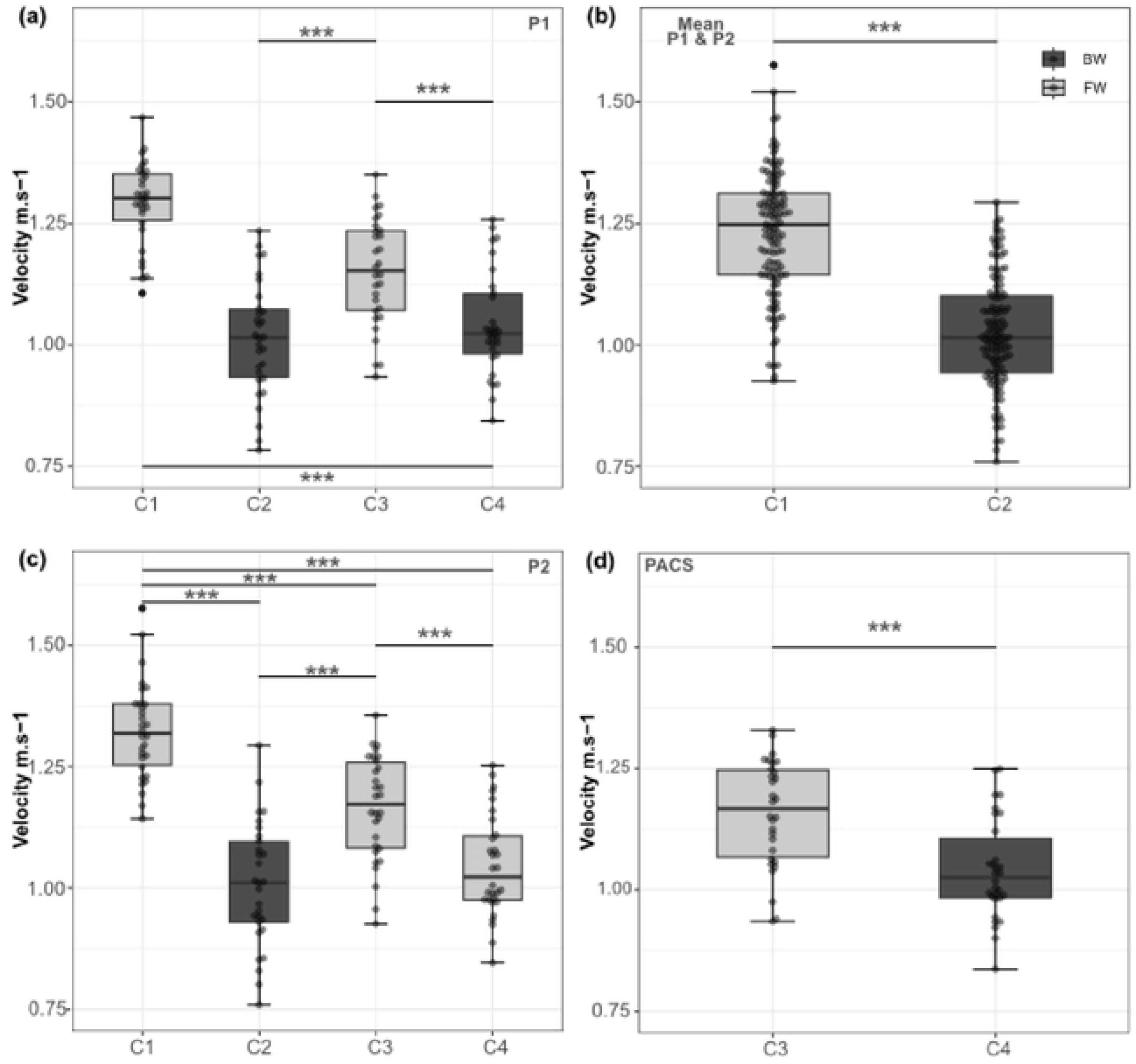
Center of Mass velocity (m.s^-1^) boxplots for participant 1 (a) and (b), participant 2 (c) and (b) and, PACS (d). for condition 1,2, 3 and, 4. Light grey boxplots represent the forward walking performance and, dark grey boxplots represent the backward walking performance(* p< 0,05; ** p<0,01 and; *** p<0,001).

### 1.9. Recovery rate

Concerning P1, statistical analysis showed a significant decrease of the RR by 38.8% from C1 to C2 and, and then by 44.3% from C3 to C4 (Figure 5.a). P2 Recovery Rate significantly decrease between C2 and C1, C3 and, C4 by 73.6%, 55.7% and, 43.9%, respectively. In addition, a P2 diminution of 10.3% from C1 to C3 and, a diminution of 17.1% from C1 to C4 were observed(Figure 5.b). Individually, the recovery rate of forward walking “C1” decreased by 40.7% compared to backward walking “C2” (Figure 5.b). In parallel, the recovery rate computed for the PACS during C3 also decreased by 27.5% compared to the C4 one (Figure 5.c).

**Figure 5:**
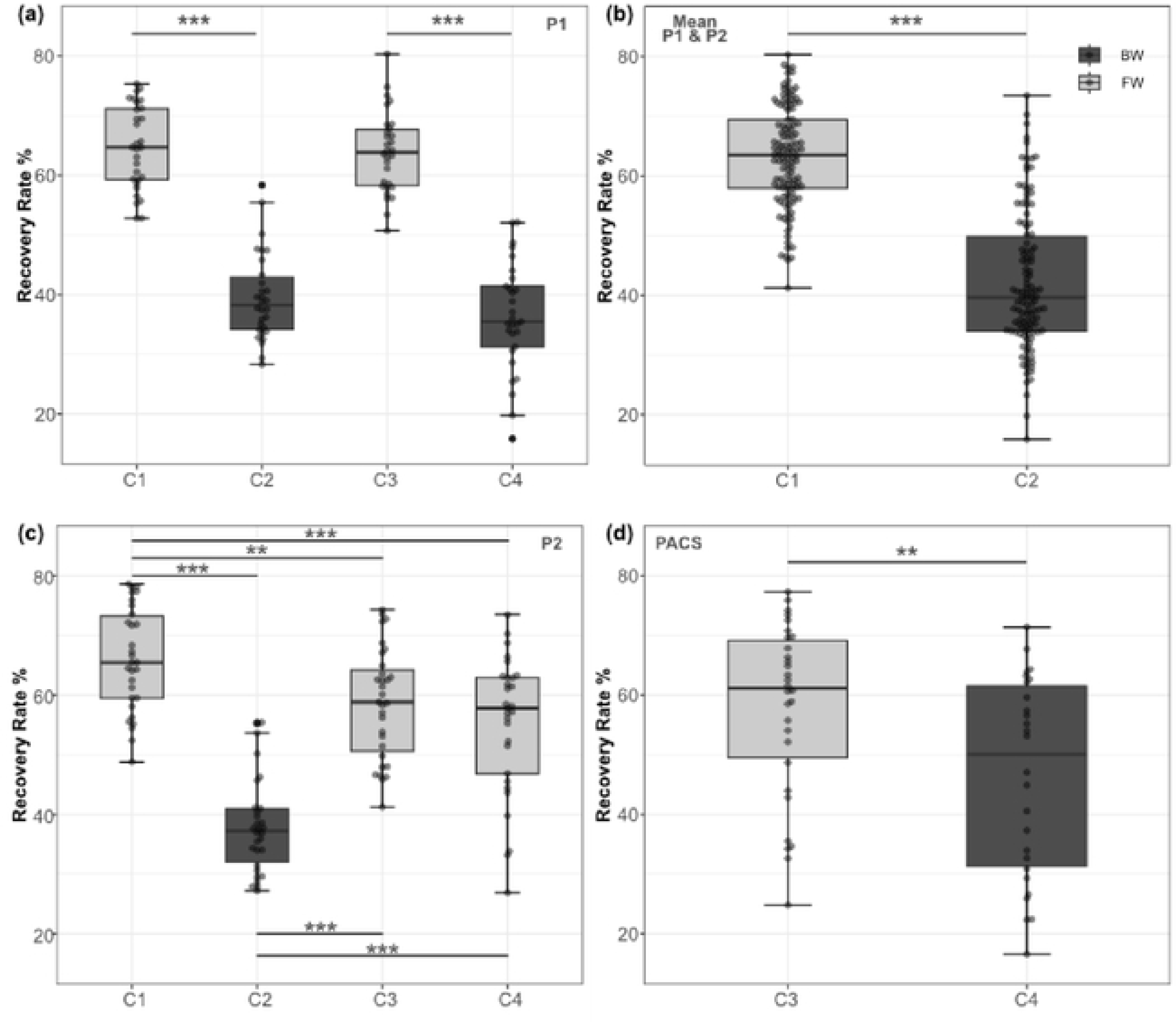
Recovery Rate (%) boxplots for participant 1 (a) and (b), participant 2 (c) and (b) and, PACS (d). for condition 1,2, 3 and, 4. Light grey boxplots represent the forward walking performance and, dark grey boxplots represent the backward walking performance (* p< 0,05; ** p<0,01 and; *** p<0,001).

### 1.10. Joint angles (Hip-Knee-Ankle)

For the participant 1, two different strategies were used depending on the walk performed (forward or backward walking). A higher flexion at the initial heel strike was observed during forward walking for participant 1. Differences were found between 40% and 80% of the gait cycle, with a decrease of 8° on average for the hip extension angle between C1 and C3 (forward walking) and between C2 and C4 (backward walking). Besides, the terminal stance occurred 8% later in backward walking compared to forward walking. The terminal stance of C1 and C3 occurred around 55% of the gait cycle, while, for C2 and C4 it occurred earlier (48% of the gait cycle). The same goes for the terminal swing, which was occurring in the early 80% of the gait cycle in the C2 and C4, whereas, it was occurring at 90% of the gait cycle in C1 and C3.

**Figure 6:**
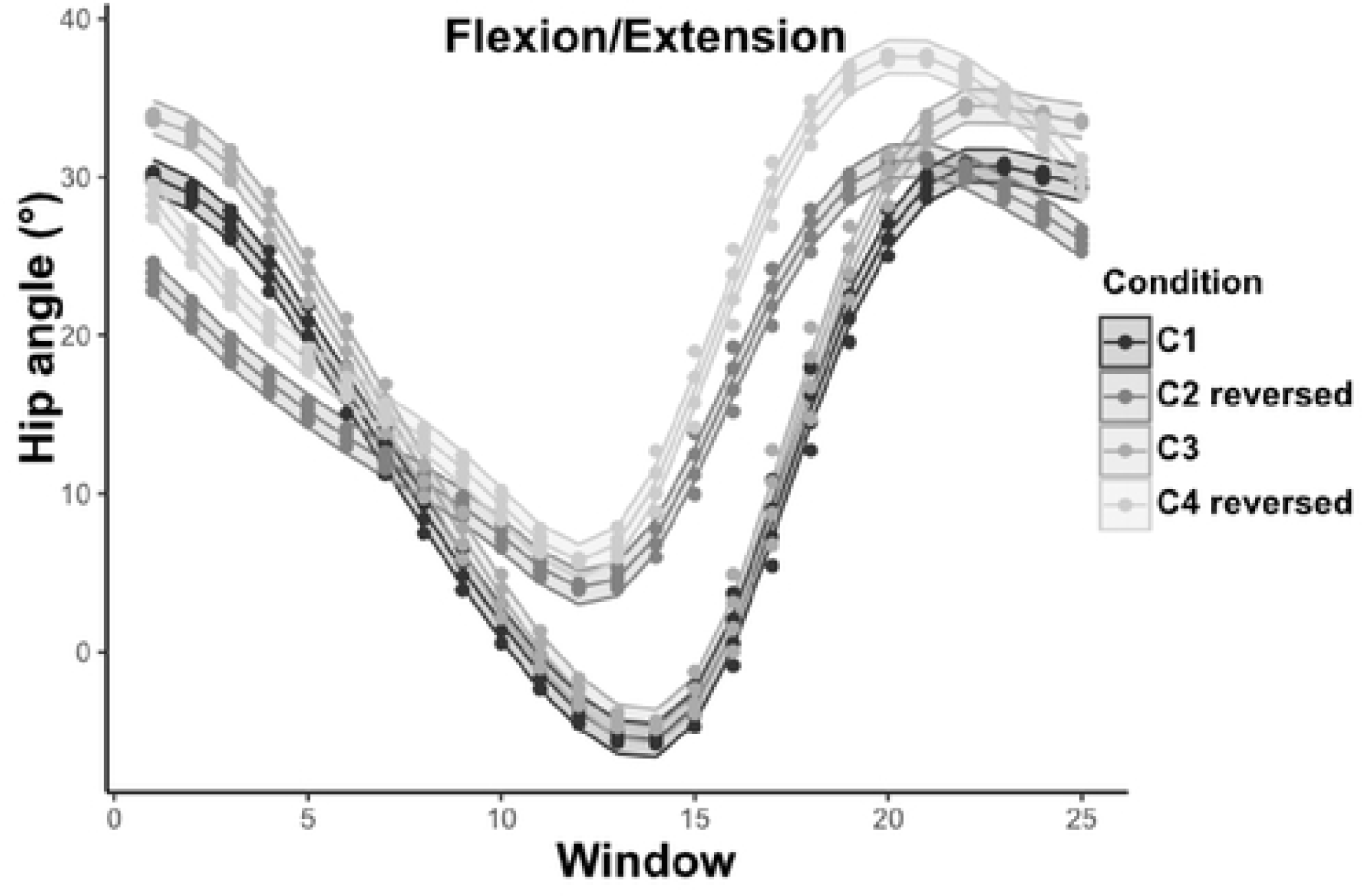
Participant 1 hip flexion/extension (°) for the condition 1, 2, 3 and, 4. The curves correspond to the 95 confidence interval of each 4% of the gait cycle. The C2 and C3 curves are reversed in time.

Concerning the participant 2, no difference was found across C1, C3 and C4. However, as observed in the participant 1, a 23.6% decrease of the initial angle at heel strike has been observed from forward to backward walking. We also found a 140.3% (9°) decrease of hip extension across forward walking (C1, C3 and, C4) and backward walking (C2) at 60% of the gait cycle.

**Figure 7:**
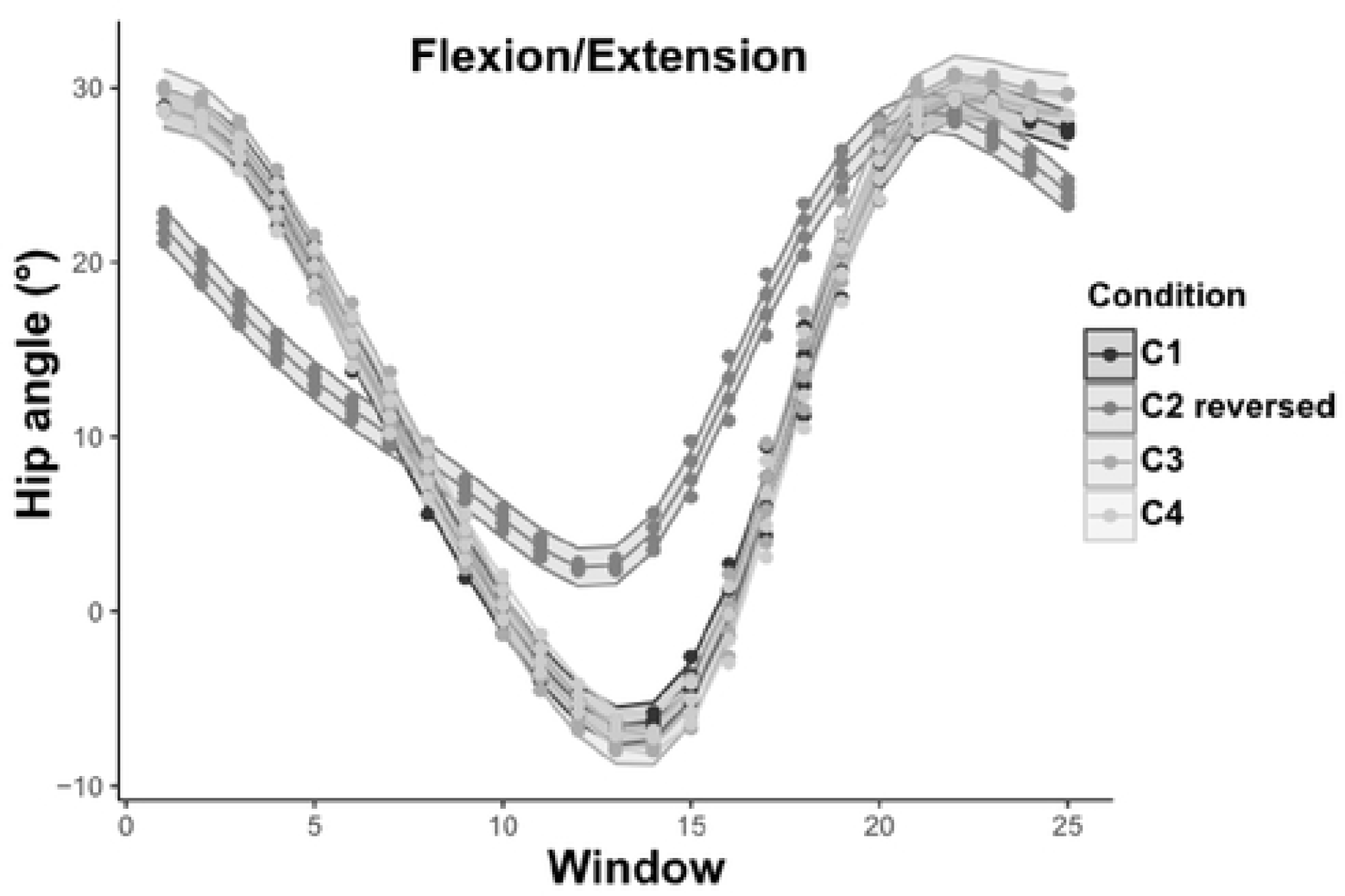
Participant 2 hip flexion/extension (°) for the condition 1, 2, 3 and, 4. The curves correspond to the 95 confidence interval of each 4% of the gait cycle. The C2 curve is reversed in time.

As for the hip, two strategies of flexion/extension occurred during the forward and backward walking. C1 and C3 represent a normal forward walking, with two flexion/extension peaks, one at 15% of the cycle and another one at 75% of the cycle. Conversely, the reversed results of backward walking, showed an erasure of the first peak associated to a direct increase of the knee flexion from 30% to 60% of the gait cycle. There was also a delay for the second peak which was occurring at 65% of the cycle. Besides, a 7° decrease of the second peak amplitude has been observed.

**Figure 8:**
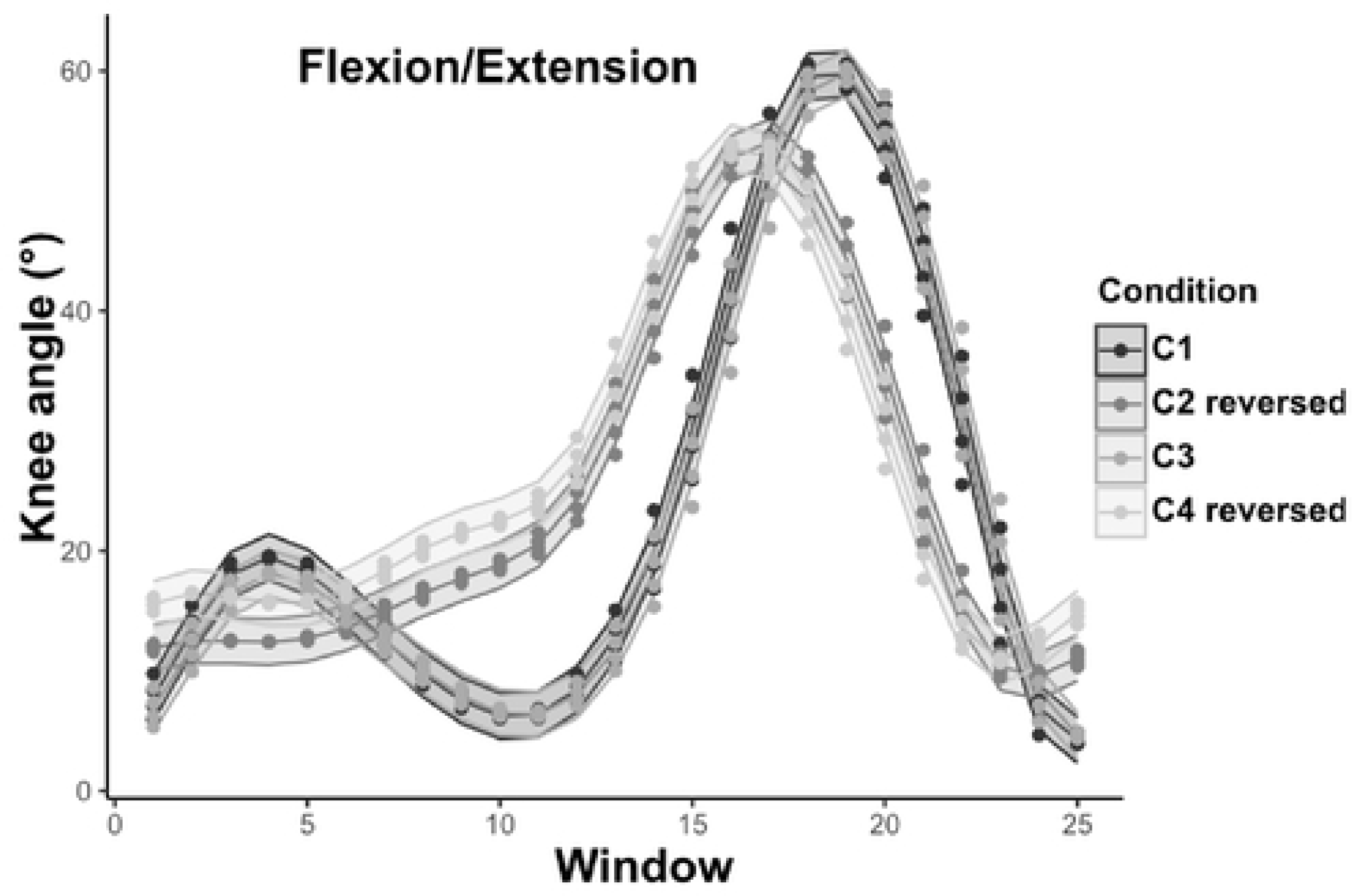
Participant 1 knee flexion/extension (°) for the condition 1, 2, 3 and, 4. The curves correspond to the 95 confidence interval of each 4% of the gait cycle. The C2 and C3 curves are reversed in time.

Participant 2 developed the same joint kinematic strategies as demonstrated at the three joints and in the three conditions (C1, C3 and, C4).

**Figure 9:**
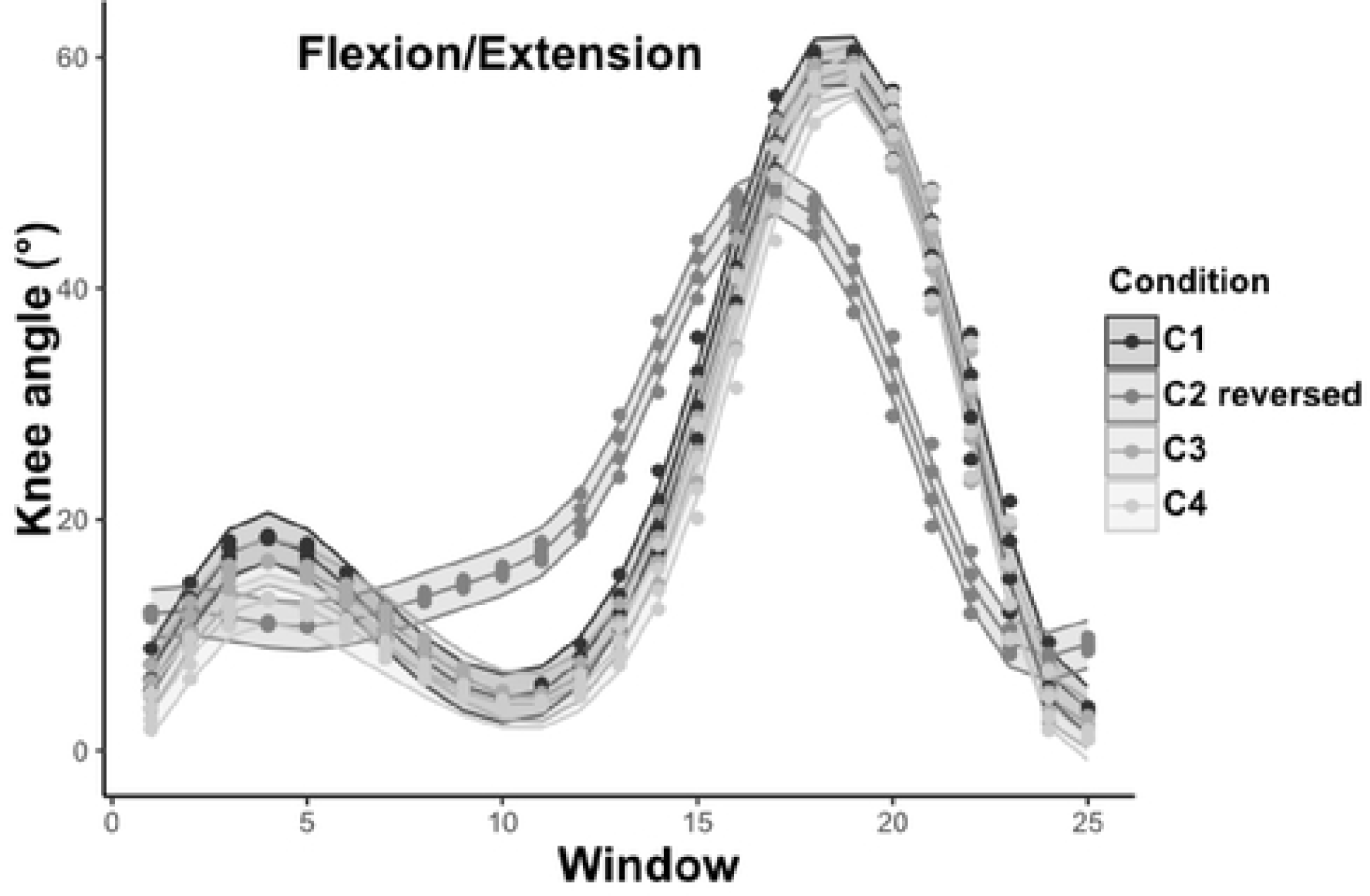
Participant 2 knee flexion/extension (°) for the condition 1, 2, 3 and, 4. The curves correspond to the 95 confidence interval of each 4% of the gait cycle. The C2 curve is reversed in time.

The results of ankle dorsi/plantar flexion reveal an increase of the dorsiflexion during C2 and C4, by 27.1% and 56.5% at 36% of the cycle, respectively. At the initial swing phase, the plantar flexion reached 12° in C3 and 9° in C1 at 65% of the cycle, while, the ankle joint reached a plantar flexion limited to approximatively 7° at 92% of the cycle. Besides, the plantar flexion was 95.3% higher in C4 collective performance than for the C2 individual performance.

**Figure 10:**
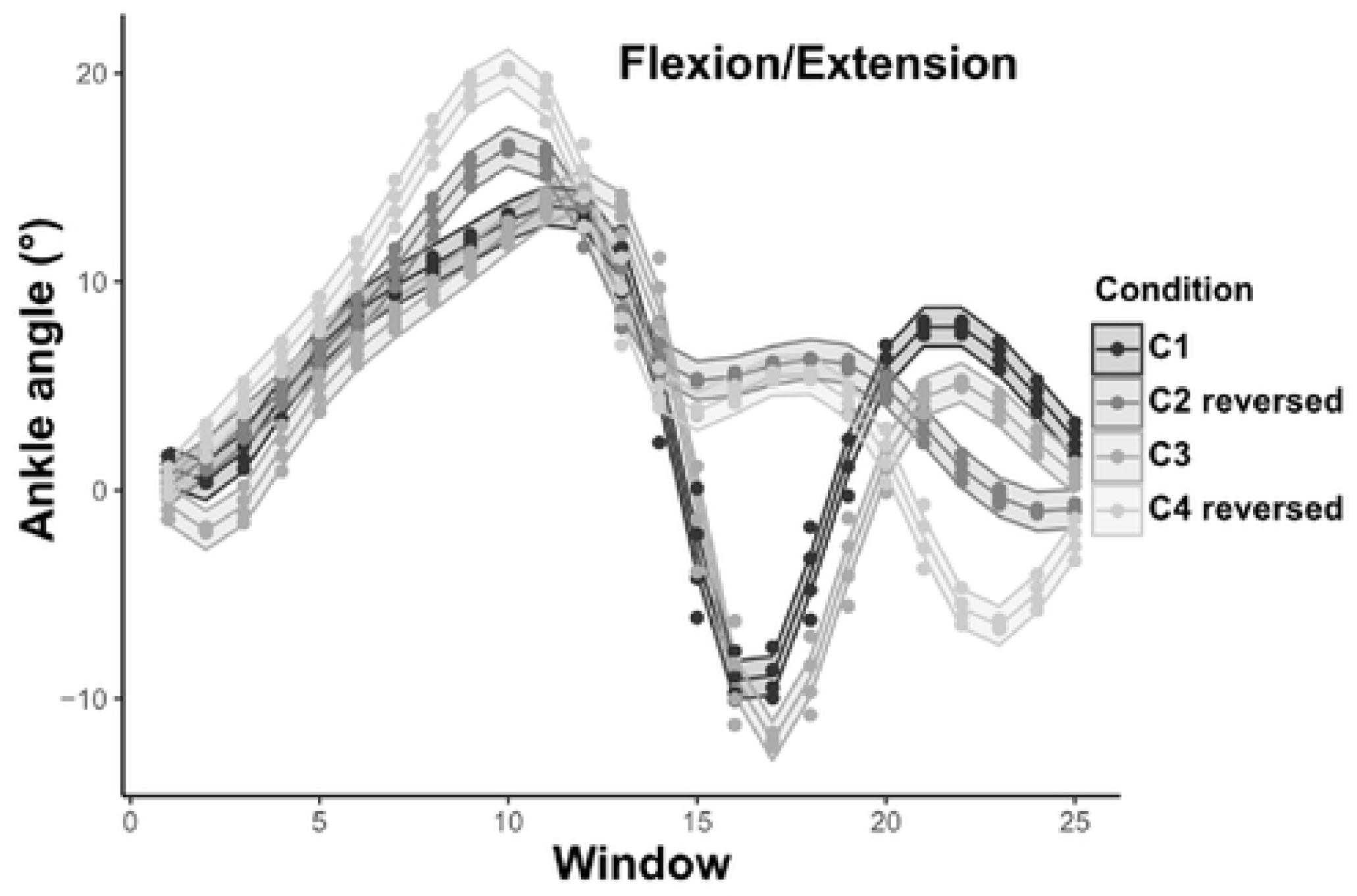
Participant 1 ankle flexion/extension (°) for the condition 1, 2, 3 and, 4. The curves correspond to the 95 confidence interval of each 4% of the gait cycle. The C2 and C3 curves are reversed in time.

The same results were observed in the participant 2, when comparing the C2 angles to the three other conditions (C1, C3 and, C4).

**Figure 11:**
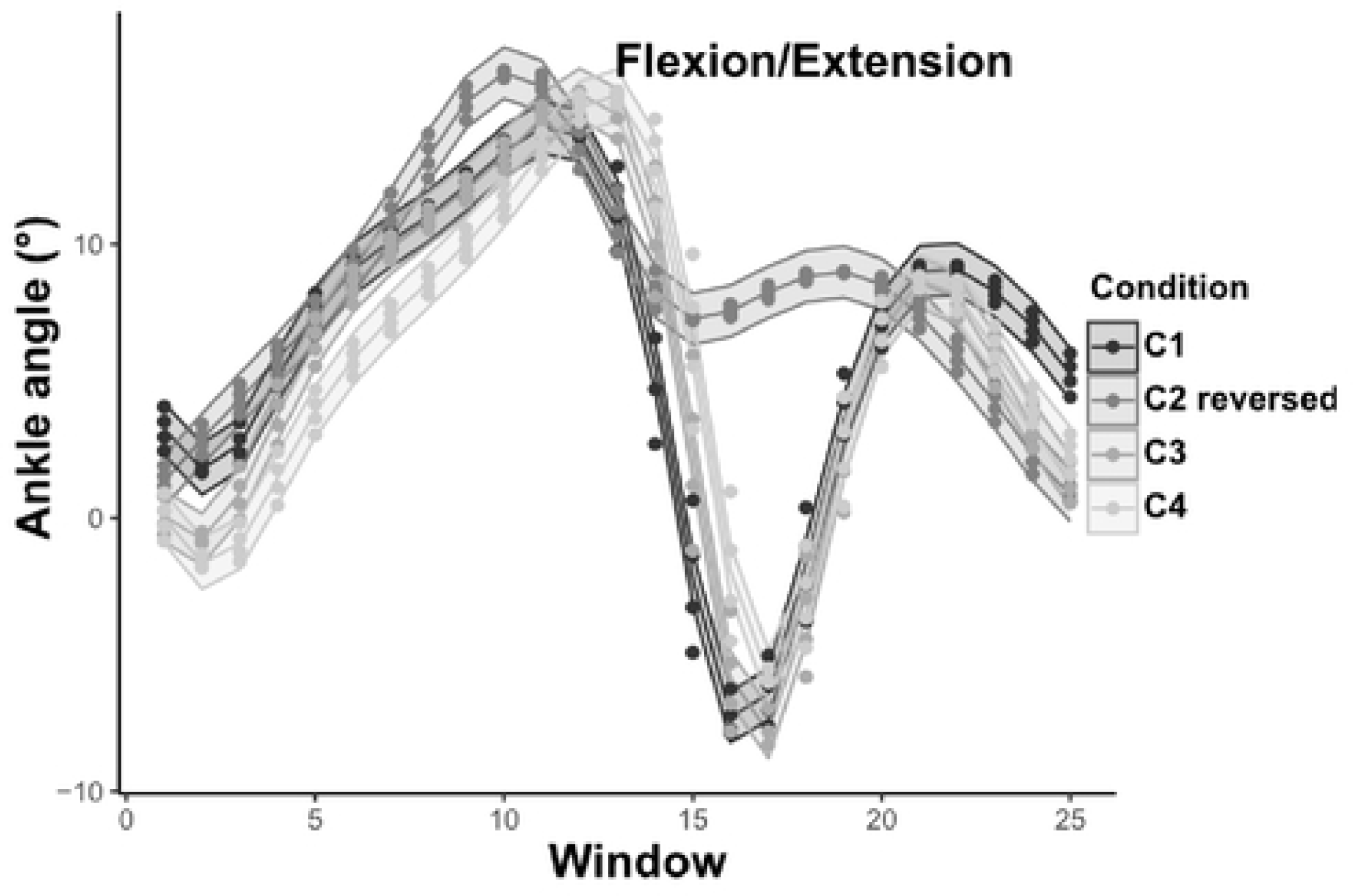
Participant 2 ankle flexion/extension (°) for the condition 1, 2, 3 and, 4. The curves correspond to the 95 confidence interval of each 4% of the gait cycle. The C2 curves is reversed in time.

## 4. Discussion

The purpose of this study investigate how a collective task such as stretcher transport influence the individual walking patterns. We used two load transport configurations with subject looking at the same direction or in the opposite direction while transporting the load, and compared the walking pattern with individual walking pattern without load. We studied the differences across forward and backward walking, compared individual walking to linked walking and, studied the impact of a backward walking on a linked walking. The results showed that the collective task modifies the spontaneous walking pattern but only/mainly for one participant.

We first compared the spatiotemporal parameters when performing a forward walking and a backward walking. Accordingly to Fritz et al., (2013) and Lee et al., (2013), a reduction of the step length and the CoM velocity during condition 2 were observed. The walking velocity is depending on the step length (JudgeRoy, Davis & Ounpuu, 1996). Unlike forward walking “C1”, backward walking “C2” is occasionally used in the human range of motion despite it alters the movement efficiency. In addition, we instructed the participants to look straight ahead during C2 meaning there was an absence of a visual feedback. Studies explained that in order to reduce their subjective instability, participants had to reduce their step length so as their speed (Grasso, Bianchi & Lacquaniti, 1998). Besides, the diminution of the average range of the hip flexion/extension induced the diminution of step length parameters (Perry, K & Davids, 1992). These results, though, did not impact the gait cycle time, which was globally around 1 second for both conditions.

The recovery rate (RR) was used as an indicator of the efficiency of the walking pattern, with the objective to assess the amount of energy transferred between the potential and the kinetic energy due to a pendulum like behavior. Globally, RR was always significantly higher when the participant performed a forward than a backward walking, whether it was an individual or collective condition. The results obtained for the individual forward “C1” and backward “C2” walking were close to the values found in the literature (Minetti & Ardigò, 2001). This same research states, that this diminution can be explained by a loss of energy, which occurs when the duty factor (the proportion of ground contact reported to the stride) approaches the 50% of the gait cycle, thus reducing the double-contact time. Concerning the collective conditions, our results were close to those obtained by Fumery et al. (2021).

The vertical Center of Mass (CoM) excursion has been widely for normal and pathological walking. However, very few described the ones occurring during a backward walking and, even less for collective transport. Our main result showed, a diminution of CoM amplitude across individual walking, forward and backward, which can be explained by the velocity decrease and the limitation of movement. This limitation occurs mainly at the terminal stance (50% of the gait cycle). In fact, for a forward walking the hip extension reaches the 10°, whereas, for a backward walking the hip extension is not present.

Most of the research done on backward walking, considered it as the reverse of a normal forward walking (Thorstensson, 1986; Winter et al., 1989; Lee et al., 2013). In this study, we compared the forward walking results to the reverse of those of backward walking. It showed, as stated by the literature, a more or less similar time-reversed pattern of flexion/extension at the hip, knee and ankle. In order to compare the range of motion evolution, Lee et al. (2013) compared only the maximum flexion/extension angles during selected crucial joint points: loading response, stance and, swing phase. In this study, we performed a sliding window of 4% to see joint angles evolution during a complete flexion/extension cycle. This statistical analysis, enable us to analyze how a backward walking (C2) impacted the gait pattern. It reveals a decrease of the range of motion and an early occurrence of some crucial points. The results showed a clear diminution of the extension for both participants when performing backward walking (C2). This result could also explain the early occurrence of the final stance, around 48% of the gait cycle. The knee flexion/extension, was also impacted by the backward walking (C2) with a decrease of the knee flexion at 80% of the gait cycle. The results obtained for the ankle correspond to those of Balasukumaran et al. (2020), who demonstrated a modification of the walking pattern kinematic. Our results confirm in some ways that the backward walking (C2) is a reversal kinematic of the forward walking (Winter et al., 1989). However, we found major modifications in joints angles amplitudes and a delay of the occurrence of some crucial time points. Those modifications can clearly explain the spatiotemporal parameters modifications as the step length decrease, velocity decrease so as the decrease of the recovery rate.

Concerning the collective transport, we recorded two conditions where the main difference was the position of the participant 1. In the C3 collective condition, both participants looked at the same direction. The results obtained showed that both participants reduced their speed when transporting the load. Multiple research showed that during individual walking, when the load weight tends to increase, the spontaneous velocity tends to decrease, due to the step length reduction (James et al., 2015). These modifications allow humans to spontaneously adopt an optimal gait and walking speed to minimize the energetic cost (Minetti & Alexander, 1997; Bode et al., 2021). Yet, in our study, we choose a negligible weight of the transported object, to focus on the effect of a physical link across the participants. This means that the observed modifications were only due to their link that induced the first step of a collaborative task.

These diminutions are amplified during the “C4”, where the participants looked at each other during transporting the stretcher-like-object. Indeed, the individual performance of P1 showed an increase of his backward velocity during “C4” compared to “C2”, when P2, on the contrary, decreased his forward velocity in “C4” compared to his performance in “C1” and “C3”. Regarding the Gait Cycle Time “CGT”, during the different conditions, P2 was the one who modified his timing depending on the performed tasks. Lanini et al.(2017), explained that global gait adaptations are mainly due to the fact that each subject tries to accommodate to the motion of the other subject which is detected by interaction forces, visual and acoustic informations. During the experiments, the participants unintentionally communicated through the interaction forces, that are considered as sensory feed-back (Zivotofsky et al. 2012).

For P1, the collective load transport did not impact his RR performance, and no difference was found across individuals and collective performance for forward and backward walking. Whereas, P2 constantly modified his behavior depending on the condition. P2 RR decreased when he performed “C3” and even more when he performed “C4”. Fumery et al.(2018b), studied paired walking of adults with intellectual disabilities and showed that when the participant is paired to a healthy individual, there is an improvement of spatiotemporal of the disabled participant and a decrease of the healthy participant pattern. In our study it seems that P2, plays an important role in the constant kinematic readjustment given that he has more environment inputs and that the backward walking disturbed P1 efficiency.

Concerning the kinematic modifications, the collective load transport “C3” slightly impacted the kinematics of participant 1 gait pattern in hip flexion/extension at the beginning and the end of the cycle. However, no other kinematic modification has been noticed for the other joint as well as for the participant 2. In general, the major modifications induced by a collective load transport are those found in spatiotemporal parameters.

The objective of the condition “C4” was to see if the backward walking performed by the participant 2 impacted the kinematic individual performance of the participant 1 as well as the collective performance. Kinematic results of participant 1 showed higher flexion values for each of the hip, knee and, ankle when comparing the individual backward performance “C2”. On the contrary, no kinematic modifications were found for the participant 2.

Regarding the diad performance, we focused on the Poly-Articulated Collective System (PACS) formed by the two participants and the load they transported. As done for the participants, we computed the PACS velocity and Recovery Rate, with the purpose to bring out how P1 placement impacted the collective performance. Both collective conditions replicate two types of collective stretcher transport. Our results showed a velocity decrease of the PACS from “C3” to “C4”. Which can be explained by the velocity individual decrease of each participant. The Recovery Rate also decreased by 10% from “C3” to “C4”. These two results join those found by Sghaier et al. (2022), who studied a collective side-by-side load transport associated to a precision constraint and, showed a decrease of the PACS velocity and Recovery Rate. The amplitude of the CoM displacement also showed a decrease from “C3” to “C4”. These results illustrate a better collective efficiency for “C3”, meaning when the two participants performed both a forward walking and looked at the same direction. This also means that the backward walking during the collective transport affects the efficiency of the collective work.

## 5. Conclusion

The present study give insight on how a collective load transport modify the individual walking performance. We first compared individual forward and backward walking. As stated by the literature we observed a kinematic time reversal of the flexion/extension angles. However, we observed major modifications in the flexion/extension amplitude and a delay of the occurrence of some crucial time points inducing a modification of some spatiotemporal parameters. Then we studied the impact of a collective load transport on the individual walking performance. When the participant looked at the same direction, we observed a slight kinematic modification of the participants. Yet, the linked task induced a decrease of spatiotemporal parameters. However, when the participants looked at each other during the carriage, we observed higher flexion values for participant 1, added to the decrease of spatiotemporal parameters. The efficiency of the collaborative task decreased when the participant looked at each other as shown the decrease of the recovery rate. This result indicates that the major component impacting the efficiency of a collective load transport was the performance of a backward walking by the participant 1. Moreover, the efficiency of the collaborative task was mainly controlled by the participant 2 with more environment information. He adjusted systematically his behavior (spatiotemporal parameter) in order to adapt to the other participant who was performing a backward walking. Future research should study the forces interaction across participants in order to understand the involvement of each participant.

## Acknowledgments

This work was supported by the Agence Nationale de la Recherche [CoBot-Projet-ANR-18-CE10-0003].

